# Extracellular vesicles, syntaxin 2 and SNAP23 collectively play a role in the receptive uterine microenvironment

**DOI:** 10.1101/2023.09.10.557108

**Authors:** Sadaf N Kalam, Samson Dowland, Louise Cole, Laura Lindsay, Christopher R Murphy

**Author notes:** Correspondence: Dr. Sadaf N. Kalam PhD, Fertility and Contraception Research Group, Faculty of Medicine and Health, Room G54, Medical Foundation Building (K25), The University of Sydney, NSW 2006, Australia., LinkedIn: www.linkedin.com/in/sadaf-kalam-b6272085.

## Abstract

Uterine luminal fluid (ULF) composition plays a major role in cell-to-cell communication between the receptive endometrium and an invading blastocyst. ULF is made up of secretions from the uterine glands and uterine epithelial cells (UEC). However, the cellular mechanisms regulating these exocytotic secretions are not yet understood. This study investigated the role of extracellular vesicles (EVs) during early pregnancy using Transmission Electron Microscopy (TEM). TEM analysis at time of fertilisation (TOF) Day 1 and at time of receptivity (TOR) Day 5.5 revealed EVs present, with an abundance at TOR. Exocytosis signalling in UECs, by SNARE proteins syntaxin 2 (syn2) and SNAP23, was also examined. Immunofluorescence microscopy showed both syn2 and SNAP23 to be present in the apical area of UECs at TOR. Western blot and immunofluorescence quantification revealed a significant increase in syn2 and SNAP23 at TOR compared to TOF. SNAP23 colocalization with apical actin showed SNAP23 was in the luminal space contributing to ULF. Overall, this data shows EVs, syn2 and SNAP23 (potential receptivity marker) are present in ULF and may together create a favourable microenvironment for blastocyst implantation.

**Summary statement:** During uterine receptivity SNAREs participate in the secretion of ULF. EVs and SNAP23 are present in the uterine luminal space during uterine receptivity. SNAP23 has the potential to be used as a receptivity marker.

## Introduction

Blastocyst implantation in the uterus involves coordinated dialogue between the blastocyst and a receptive endometrium (Carson et al., 2000; Lessey, 2000; Murphy, 2004). The endometrium and specifically the luminal uterine epithelial cells (UECs) undergo defined morphological changes to establish uterine receptivity. This involves the loss of regular microvilli and non-adhesive proteins from the apical plasma membrane and reorganisation of microtubules and the actin terminal web (Aplin, 1997; Kalam et al., 2018; Moore et al., 2016; Murphy, 2004). Instead, at the time of uterine receptivity, the apical cell surface has adhesive proteins, pinopods, and increases in plasma membrane cholesterol, porosomes and in intramembranous particles (Kalam et al., 2022; Murphy, 1993; Murphy, 1994; Murphy, 2004; Murphy and Dwarte, 1987; Murphy and Martin, 1985; Singh and Aplin, 2009). Additionally, fluid changes in the luminal microenvironment also play a major role in blastocyst implantation (Aghajanova et al., 2008; Erranz et al., 2004; Glasser et al., 1987; Ng et al., 2013; Salamonsen et al., 2016).

Uterine luminal fluid (ULF), once considered simply the medium for sperm and embryo transport, is now thought to play a major part in co-ordinating cell to cell communication between a receptive endometrium and an invading blastocyst (Zhang et al., 2017). It is also known that disruption of the volume or molecular content of uterine fluid homeostasis can cause abnormal embryo implantation and pregnancy loss (Lu et al., 2013; Zhang et al., 2015). Due to its extracellular location, it can be readily acquired and studied non-invasively. Thus, understanding the composition of ULF is of great value for determining the key requirements for uterine receptivity and thus has potential for improving artificial reproductive technologies (ART), such as via identification of biomarkers of the time of uterine receptivity.

Extracellular vesicles (EVs) have been observed in ULF of the sheep uterus and in the human reproductive tract, where the uterus is thought to be one of the possible source of these EVs (Burns et al., 2014; Tannetta et al., 2014). In cell culture exosomes have been reported to be secreted by uterine endometrial cancer cells (ECC) (Franchi et al., 2016; Ng et al., 2013; Nguyen et al., 2016). Thus, this study investigated EVs that are part of ULF secretions during early pregnancy.

EVs are categorised into three groups depending on diameter, surface markers and mode of release from host cells. These include exosomes (50-100nm), microvesicles (MVs) (100nm- 1μm) and apoptotic vesicles (50nm-5μm) (Crescitelli et al., 2013; Denzer et al., 2000; EL Andaloussi et al., 2013) as measured by various isolation and characterisation techniques such as differential centrifugation, density gradient centrifugation, size exclusion chromatography, polyethylene glycol (PEG) precipitation, precipitation with chemicals, immunoprecipitation, ultrafiltration, microfluidic devices, bronchoalveolar lavage fluid (BALF), dynamic light scattering (DLS), nanoparticle tracking analysis (NTA), nanoscale flow cytometry (nanoFACS) and Transmission Electron Microscopy (TEM) (Carnino et al., 2019). Other reported EVs such as oncosomes (100nm to 10μm in diameter) found in various other tissues fall in between or outside of these categories and are therefore instead defined by the tissue or disease origin (Di Vizio et al., 2012; Morello et al., 2013; Ronquist and Brody, 1985). In secretory cells, exosomes are formed in endosomal compartments called multivesicular bodies (MVBs) and fusion of MVBs to the plasma membrane releases smaller vesicles which are the exosomes (Bobrie et al., 2011). MVs, on the other hand, are regions of plasma membrane that bud out and are pinched off as EVs (Antonyak and Cerione, 2014; Cocucci et al., 2009). Both of these types of EVs carry and transfer regulatory molecules such as microRNAs, proteins and lipids that may mediate intercellular communication (Bebelman et al., 2018; Marca and Fierabracci, 2017; Raposo and Stoorvogel, 2013).

ULF is generated by combination of secretion form the uterine glands and UECs (Filant and Spencer, 2014; Toner and Adler, 1985), however mechanisms regulating these secretions are not yet understood. Soluble NSF Attachment Proteins (SNAPs) and SNAP receptors (SNARE) proteins are a large superfamily of proteins that mediate vesicle fusion. SNARE proteins along with Ca^2+^ signalling regulate exocytosis, thereby controlling vesicular release (Jahn and Südhof, 1999; Malmersjö et al., 2016). Secretion involves the release of vesicular content with fusion of vesicle membrane to its target plasma membrane. This requires a vesicle SNARE (v-SNARE) such as vesicle associated membrane protein (VAMP) located on a vesicle membrane and two target SNAREs (t-SNARE), a syntaxin and a SNAP23/25 located in the target plasma membrane to form a SNARE core complex (Söllner et al., 1993; Sutton et al., 1998). This complex will traffic and regulate apical luminal secretions (Hong, 2005; Nichols et al., 1997). In previous studies examining secretions form polarised epithelial cells in lung and pancreas tissue, syntaxin 2 and SNAP23 have been shown to control apical secretions (Abonyo et al., 2004; Imai et al., 2003). The present study examined the role of these proteins in generating apical secretions in UECs and contributing to the production of ULF during early pregnancy. More specifically, this study identified and characterised the various types of exosomes and MVs that are present in the uterine luminal space at the time of fertilisation and time of uterine receptivity in the rat uterus. We further investigated the roles of syntaxin 2 and SNAP23 in UEC secretion by examining the localisation and abundance of these proteins in UECs during early pregnancy. This research provides insights to the mechanisms involved in the production of ULF, which makes a significant contribution to the uterine microenvironment and development of uterine receptivity. Overall, this study inspected the secretory contents and identified SNAP23 as a key component in ULF during early pregnancy in the rat.

## Materials and methods

### Animals and mating

This study used female virgin Wistar rats aged 10-12 weeks and all procedures were approved by The University of Sydney Animal Ethics Committee. Rats were housed in plastic cages at 21°C under a 12 hour light-dark cycle and were provided with free access to food and water. Pro-oestrus female rats were mated overnight with males of proven fertility. The presence of sperm in a vaginal smear the following morning indicated successful mating, and this was designated day 1 of pregnancy. Uterine tissues were collected from five rats each on days 1, 3.5, 5.5, 6 and 7 of pregnancy for immunofluorescence and western blot analysis. Further uterine tissue was collected from four rats each on days 1, 5.5 and 6 for transmission electron microscopy (TEM).

Rats were administered 20 mg/kg of sodium pentobarbitone (Vibac Animal Health, NSW, Australia) intraperitoneally and the uterine horns were collected under deep anaesthesia, before euthanasia. The uterine horns were then used for immunofluorescence microscopy, western blotting, and TEM.

### Transmission Electron Microscopy

Uteri were cut into 5mm pieces and fixed in Karnovsky’s fixative [2.5% glutaraldehyde (ProSciTech, Australia), 2% paraformaldehyde (ProSciTech) in 0.1 M Sorenson’s phosphate buffer (PB, pH 7.4)] for 45 mins at RT. The tissue was further cut into 0.5-1 mm slices under a droplet of fixative and returned to fresh fixative for another 45 mins. The tissue was washed twice for 5 mins in 0.1 M PB then washed again with 0.1 M maleate buffer (MB), twice 5 min. Next the tissue was post fixed with 1% tannic acid in 0.1 M MB for 40 mins. Tissue was rinsed in 0.1M MB twice for 5 min, then dH20 twice for 5 min and further fixed with 1% uranyl acetate in dH20 for 1 hour. Tissue was rinsed again in dH20 three times for 5 min and dehydrated with a graded series of ethanol then infiltrated with Spurr’s resin (SPI supplies, Leicestershire, England, UK). Samples were embedded in fresh Spurr’s resin in BEEM^®^ capsules (ProSciTech) and polymerised at 60 °C for 24 h. Two blocks per animal were cut using a Leica Ultracut S ultramicrotome and sections 60-70 nm thick were mounted onto 400-mesh copper grids. Sections were post-stained with 2% uranyl acetate in dH20 for 10 mins and then with Reynold’s lead citrate for 10 mins. Sections were examined in a Jeol 1400 TEM (Jeol Ltd., Japan) at 100 kV.

### Immunofluorescence microscopy of syntaxin 2 and SNAP23

Uterine horns (5mm pieces) were coated with OCT (Tissue Tek, CA, USA), snap frozen in supercooled isopentane, and stored under liquid nitrogen until use. Frozen sections (7µm) were cut using a Leica CM 3050 cryostat (Leica, Heerbrugg, Switzerland) and air-dried onto gelatine-chrome alum-coated slides.

Sections for syntaxin 2 staining were fixed in 4% formaldehyde in 0.1 M phosphate buffer while those for SNAP23 were fixed in 75% methanol in phosphate buffered saline (PBS) for 10mins at room temperature (RT). All sections were subsequently washed with PBS and blocked with 1% bovine serum albumin (BSA; Sigma Aldrich, MO, USA) in PBS for 30 min at RT. Sections were incubated overnight at 4°C with rabbit monoclonal anti-syntaxin 2 antibody (7.8 µg/ml; Abcam, Cambridge, England, UK: ab170852) or goat polyclonal anti- SNAP23 antibody (1.25 µg/ml; Abcam, ab166808), diluted in 1% PBS/BSA. Concurrently, control sections were incubated with non-immune mouse or goat IgG (Sigma Aldrich) at the same concentration as the primary antibody. Sections were washed in PBS and incubated with FITC-conjugated goat anti-mouse IgG (3µg/mL; Jackson ImmunoResearch, PA, USA) or Alexa Fluor 488 conjugated donkey anti-goat IgG (0.02 µg/ mL; Life Technologies Australia Pty Ltd, Mulgrave, VIC, Australia) for 30 minutes at RT. Sections were washed in PBS and mounted onto glass slides with Vectashield containing DAPI (Vector Laboratories, CA, USA) and coverslipped (No. 1 thickness).

Zeiss AxioImager M2 Microscope (Carl Zeiss, Australasia) was used to image the sections and images were acquired with the use of Zeiss AxioCam HR digital monochrome CCD camera (Carl Zeiss) and ZEN 2013 (Blue edition) software (Carl Zeiss).

The camera exposure time was set on the brightest sample per protein and the same parameters were used to image all other samples. These images were then used to quantify the fluorescence intensity.

### Image analysis

Intensity measurements were made using FIJI (Fiji is just ImageJ). Uterine epithelial cells were selected for intensity measurements. Cell counts were obtained from the DAPI channel, by thresholding the nuclei and a particle size limit were set to eliminate any peripherally cut cells in the section. The FITC or Alexa 488 channel with the protein staining was thresholded at the same range for all images and a gray scale value was obtained for each image. These were then standardised to the number of nuclei to obtain a measurement of fluorescence intensity per cell. Three randomly selected high magnification images per rat were analysed and averaged to provide an average intensity per set. Unpaired two-tailed Student’s t-test analysis were performed and P<0.05 was determined to be significant. All graphs were generated using GraphPad Prism Software (Version 7.02, GraphPad Software, Inc., CA, USA) and error bars represent mean ± SEM.

### Co-localisation analysis

For co-localisation analysis of SNAP23 and Phalloidin-staining, frozen sections of day 5.5 pregnant uteri were fixed in 50/50 absolute methanol/PFA for 10 mins at RT. Then sections were washed with PBS and blocked with 1% BSA (Sigma Aldrich) in PBS for 30 min at RT. Sections were then incubated overnight at 4°C with goat polyclonal SNAP23 (1.25 µg/ml; Abcam, ab166808), diluted in 1% PBS/BSA. Sections were washed in PBS and incubated with Alexa Fluor 488 donkey anti-goat IgG (0.02 µg/mL; Life Technologies) for 30 minutes at RT. Concurrently, control sections were incubated with non-immune IgG (Sigma Aldrich) at the same concentration as primary antibody. Sections were washed in PBS then incubated with tetramethylrhodamine isothiocyanate (TRITC)-conjugated Phalloidin (Sigma-Aldrich) diluted to 0.5 µg/mL in 1% BSA/PBS for 1 hr to stain filamentous actin (F-actin). Sections were then washed with PBS and mounted with Vectashield with DAPI (Vector Laboratories) and coverslipped (No. 1 thickness). Alongside, SNAP23-only and Phalloidin-only sections were prepared as controls. Confocal images were taken using a Zeiss LSM 800 confocal microscope (Carl Zeiss). Pearson co-localisation coefficient (PCC) was calculated using the co-localisation module in the ZEN 2 software (Carl Zeiss). PCC close to one indicates protein co-localisation.

### Isolation of rat uterine luminal epithelial cells

UECs were isolated as previously described (Kaneko et al., 2008) and immediately placed into lysis buffer (50 mM Tris–HCl, pH 7.5, 1 mM EDTA, 150 mM NaCl, 0.1 % SDS, 0.5 % Deoxycholic acid, 1 % Igepal and 1% protease inhibitor cocktail; Sigma Mammalian Cell lysis kit, Sigma Aldrich) with 10% PhosSTOP phosphatase inhibitor (Roche, NSW, Australia). The isolated cells were homogenised using a 1 ml syringe (Livingstone International, Rosebery, NSW, Australia) and centrifuged at 8,000 **g** at 4 °C for 3 min. The supernatant was collected and frozen immediately in liquid nitrogen and stored at −80 °C until use for western blotting.

### Western Blot analysis

Protein concentrations were determined using the BCA protein assay (Micro BCA^TM^ protein assay kit; Thermo Fisher Scientific, MA, USA) and CLARIOstar microplate reader (BMG labtech Durham, NC, USA) according to the manufacturer’s instructions. For detecting syntaxin 2, protein samples (20 µg) and sample buffer (8 % glycerol, 50 mM Tris–HCl, pH 6.8, 1.6 % SDS, 0.024 % bromophenol blue, 4 % dithiotheitol (DTT)) were heated at 95 °C for 10 min prior to loading. For detecting SNAP23, protein samples (20 µg) and sample buffer (8 % glycerol, 50 mM Tris–HCl, pH 6.8, 1.6 % SDS, 0.024 % bromophenol blue, 4 % β-mercaptoethanol) were heated at 95 °C for 5 min prior to loading. Protein was separated on 12 % FastCast SDS-polyacrylamide gel (Bio-Rad Laboratories, Hercules, CA, USA) through electrophoresis at 200 V for 40 min and transferred to a polyvinylidene dilfluoride (PVDF) membrane (Immunobilon™ transfer membrane; Millipore, Bedford, MA, USA) at 100 V for 90 mins. Membranes stained with syntaxin 2 were blocked in 1% skim milk in TBS-t (10 mM Tris–HCl, pH 7.4, 150 mM NaCl, 0.05 % Tween 20) whilst those stained with SNAP23 were blocked in 5% skim milk for 1 h RT with constant agitation. Membranes were then incubated with syntaxin 2 (1.95 µg/ml, Abcam, ab170852) or anti-SNAP23 (0.25 µg/ml, Abcam, ab166808) primary antibodies diluted in 1 % skim milk in TBS-t overnight at 4 °C on a rocking platform. The membranes were washed in TBS-t and subsequently incubated with goat anti-rabbit IgG horseradish peroxidase-conjugated secondary antibody (0.5 µg/ml; Dako, VIC, Australia) or rabbit anti-Goat IgG horseradish peroxidase-conjugated secondary antibody (0.125 µg/ml; Dako) for 2h at RT with constant agitation. Protein bands were detected with Immobilon Western HRP Substrate (Merck Millipore) and images captured with a CCD camera and the Bio-Rad ChemiDoc MP System (Bio-Rad). Membranes were then incubated in stripping buffer [62.5 mM Tris–HCl (pH6.7), 2 % SDS and 100 mM β-mercaptoethanol] at 60 °C for 45 min and reprobed with mouse monoclonal anti-β-actin antibody (0.4 μg/ml; Sigma Aldrich) overnight at 4°C and HRP-conjugated goat anti-mouse IgG (0.2 μg/mL; GE Healthcare) for 2hr at RT, to ensure equal loading.

### Densitometry analysis

Protein band intensities were quantified with Bio-Rad Image Lab 4.0 software (Bio-Rad) using the Volume Analysis Tool with local background subtraction from the and normalised to β-actin band intensities from the same lane. Statistical analysis was performed on normalised intensities with GraphPad Prism Software (Version 7.02, GraphPad Software). Changes in quantity from day 1, 3.5, 5.5, 6 and 7 were analysed using ordinary one-way ANOVA. For multiple comparisons Turkey’s post hoc test was applied (reporting multiplicity-adjusted P-values) to determine which pairs of means were significantly different.

## Results

### EVs are present in the uterine lumen during early pregnancy

EVs are present on day 1, 5.5 and day 6 of pregnancy in the luminal cavity of the uterus when examined by TEM (Fig. 1, 2; Table 1). EVs measured to be 100 nm or smaller in diameter were classified as exosomes. Exosomes were observed in the lumen on day 1 of pregnancy (Fig. 1A). These exosomes exhibited two different types of membranes, classified as exosome type 1 and type 2. Exosome type 1 ranged from 20 nm to 100 nm in diameter with a membrane thickness up to 25 nm (Table 1). Exosome type 2 ranged from 10 nm to 60 nm in diameter with a membrane thickness up to 10 nm (Table 1). Other EVs with a diameter higher than 100nm were classified as MVs. MVs were observed in the lumen on day 1, 5.5 and 6 of pregnancy with three different types of membranes (Fig. 1; Table 1). They were classified as MVs type 1, 2, and 3. MVs generally range from 100nm to 1000nm. However, the MVs observed on day 1, 5.5 and 6 ranged from 100nm to 4100nm. Type 1 MVs ranged from 100 nm to 2500 nm in diameter with a membrane thickness up to 25 nm. MV type 2 ranged from 100 nm to 4100 nm in diameter with a membrane thickness up to 150 nm. MV type 3 ranged from 100 nm to 1000 nm in diameter with a membrane thickness up to 300 nm (Table 1). On day 5.5 numerous EVs were observed in the uterine lumen (Fig. 2). These EVs were to be associated with pinopods. Numerous EVs were seen to cluster and fuse together regardless of size and membrane composition on day 1, 5.5 and 6 (Fig. 1, 2). Additionally, multivesicular bodies as EVs were also found in the uterine lumen on day 1 and 6 (S Fig. 1A, B).

**Figure 1:**
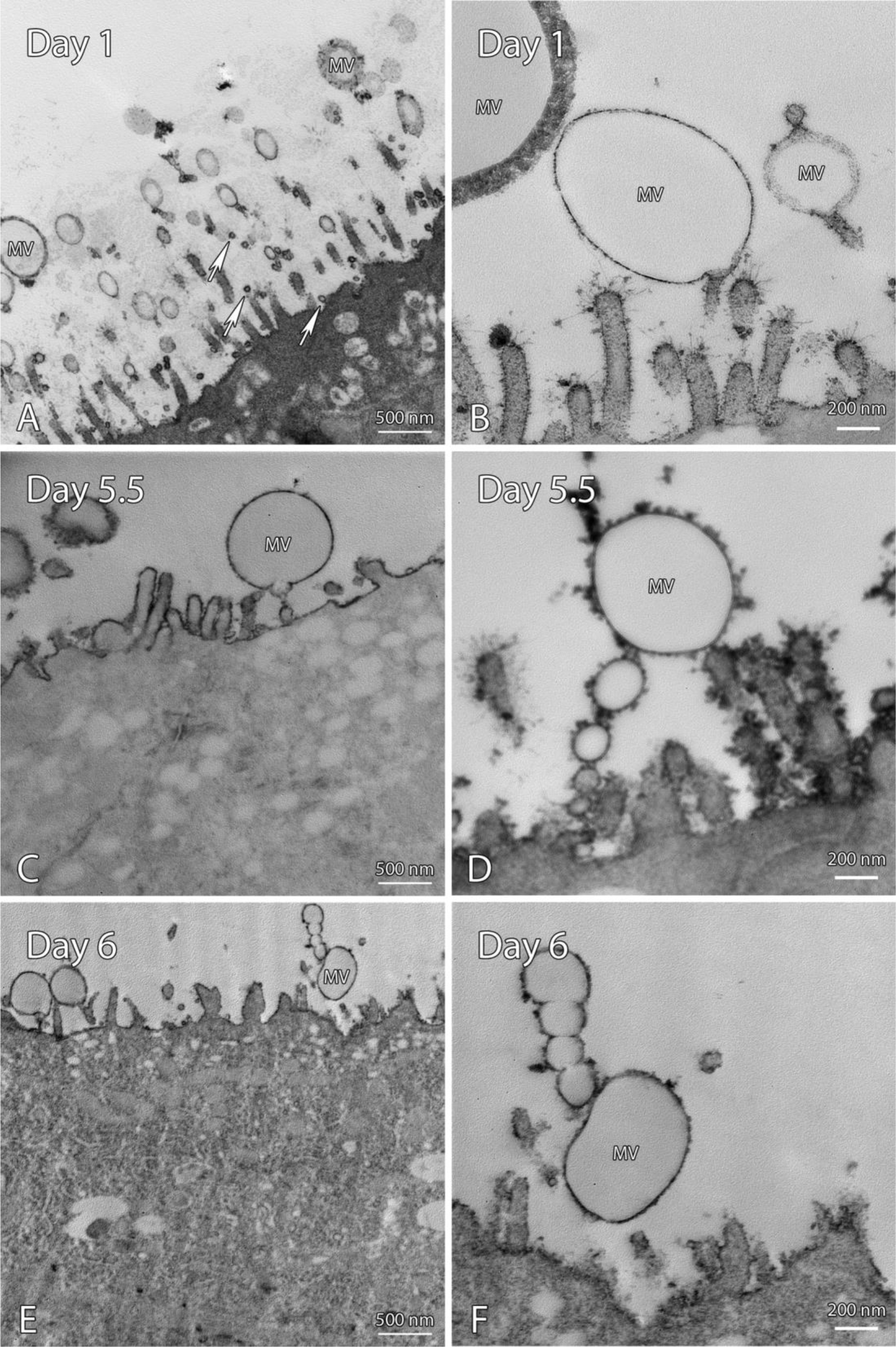
Transmission electron microscopy of extracellular vesicles in UECs on day 1, 5.5 and 6 of early pregnancy Extracellular vesicles were present in day 1, 5.5 and 6 of pregnancy. (A, B) Day 1 of pregnancy found exosomes (arrows) and microvesicles (MV) within the luminal space of the uterus. (C, D, E, F) Microvesicles were found in day 5.5 and 6 of pregnancy budding off from the UECs. All scale bars are 200nm.

**Figure 2:**
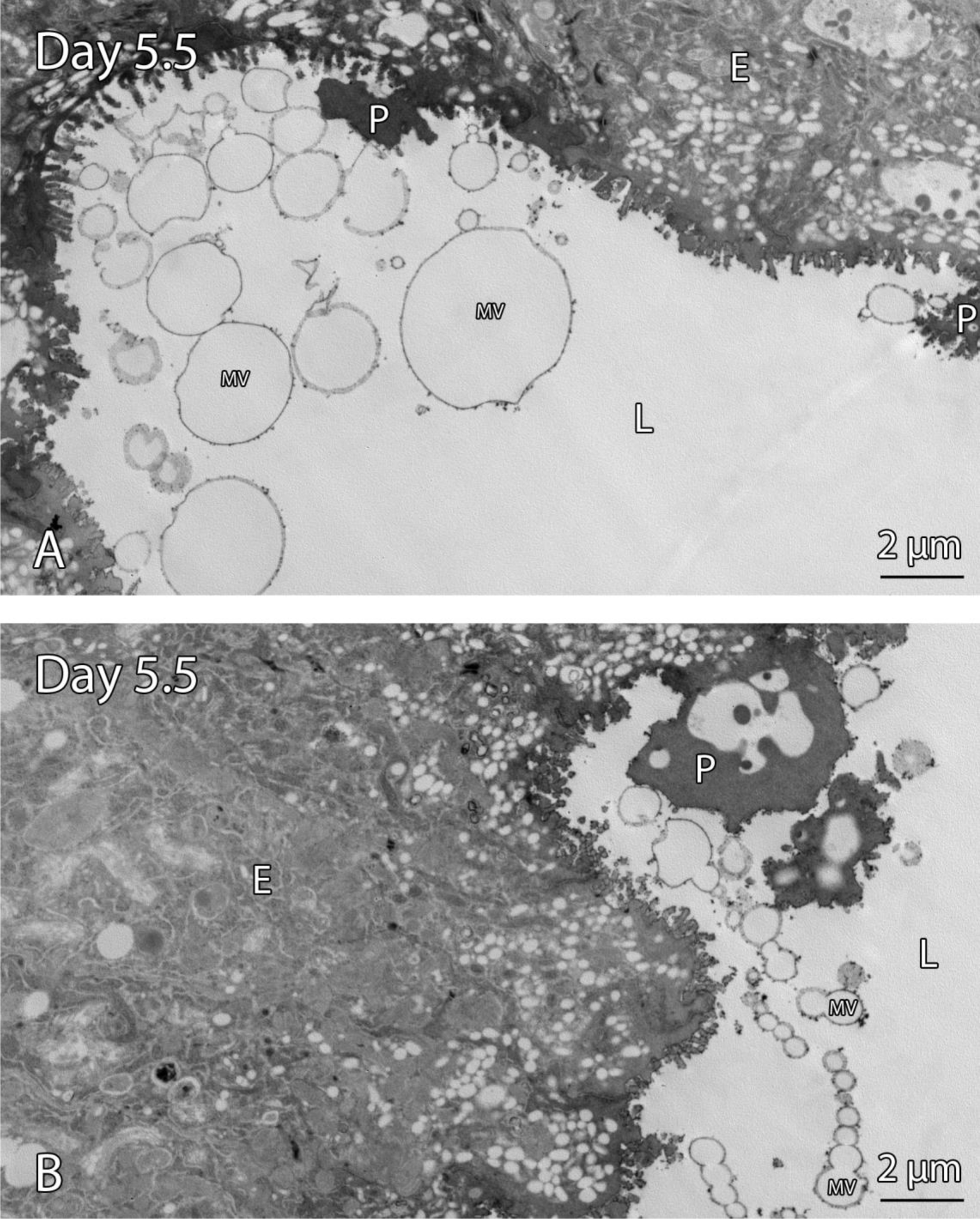
Transmission electron microscopy of extracellular vesicles in UECs on day 5.5 of early pregnancy (A, B) A large number of microvesicles (MV) with various sizes and membrane compositions are secreted into the luminal space on day 5.5 of pregnancy. These microvesicles are found near pinopods (P) and appear to aggregate and accumulate together. All scale bars are 200nm.

**Table 1:**
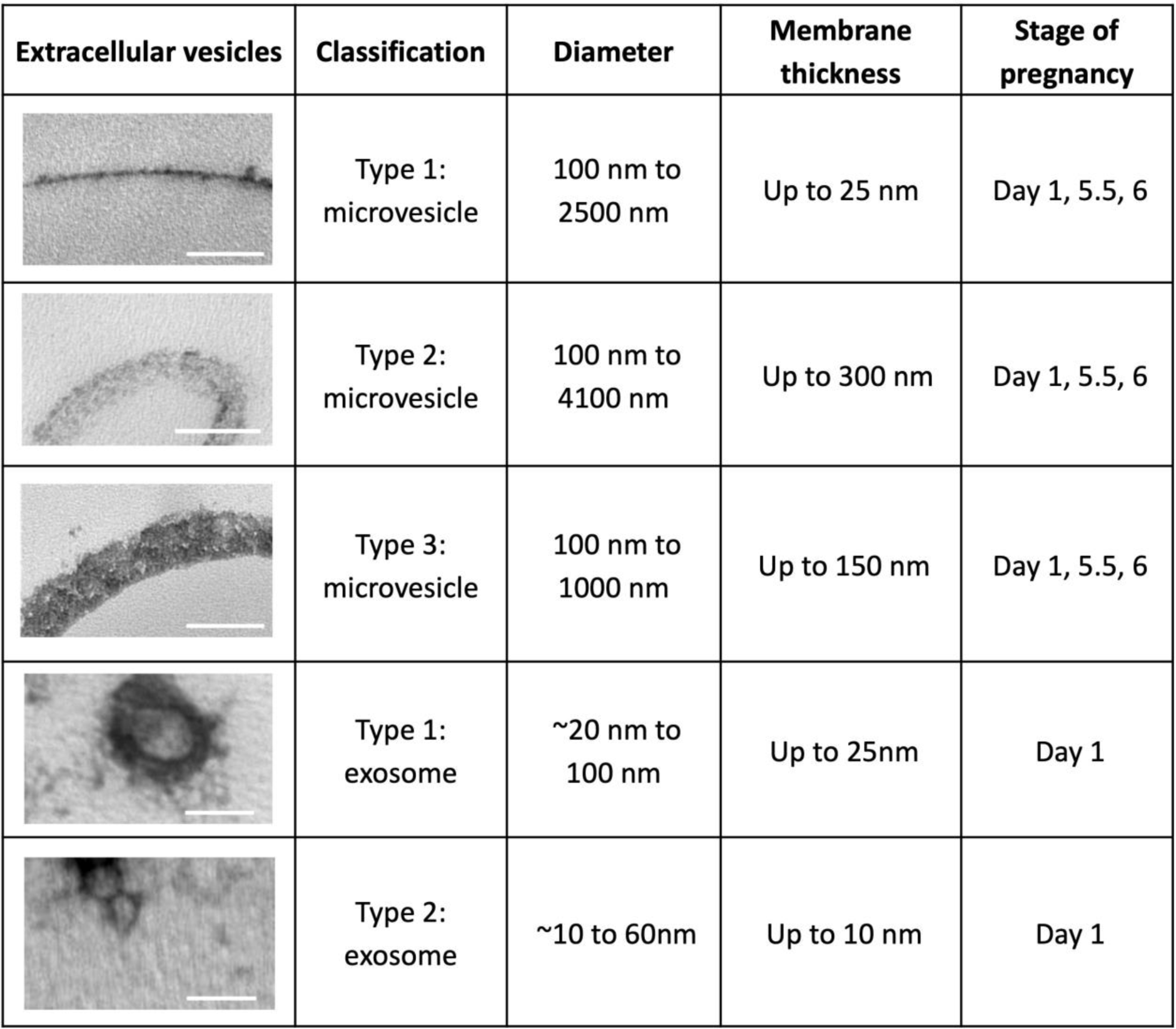
Extracellular vesicle classification found in day 1, 5.5 and 6 of pregnancy. Scale bars for MV are 200 nm and exosomes are 50 nm.

| Extracellular vesicles | Classification | Diameter | Membrane thickness | Stage of pregnancy |
| --- | --- | --- | --- | --- |
|  | Type 1: microvesicle | 100 nm to 2500 nm | Up to 25 nm | Day 1, 5.5, 6 |
|  | Type 2: microvesicle | 100 nm to 4100 nm | Up to 300 nm | Day 1, 5.5, 6 |
|  | Type 3: microvesicle | 100 nm to 1000 nm | Up to 150 nm | Day 1, 5.5, 6 |
|  | Type 1: exosome | ~20 nm to 100 nm | Up to 25nm | Day 1 |
|  | Type 2: exosome | ~10 to 60nm | Up to 10 nm | Day 1 |

### Syntaxin 2 is present in UECs during early pregnancy

Indirect immunofluorescence labelling and florescence microscopy revealed syntaxin 2 is present in UECs during early pregnancy in the rat uterus (Fig. 3). On day 1, 3.5 and 7 of pregnancy syntaxin 2 is found cytoplasmically throughout the cell (Fig. 3A, B, E). On day 5.5 and 6, the time of receptivity, syntaxin 2 is localised in the cytoplasm and apically in UECs (Fig. 3C, D). Non-immune controls performed with all immunofluorescence protocols showed no staining in UECs. A representative image of day 5.5 non-immune controls is shown (Fig 3F). Measurements of fluorescence intensity in image data taken from days 1 and 5.5 found significantly more syntaxin 2 per epithelial cell on day 5.5 compared to day 1 of pregnancy; *p <0.05; n=5 (Fig. 3G).

**Figure 3:**
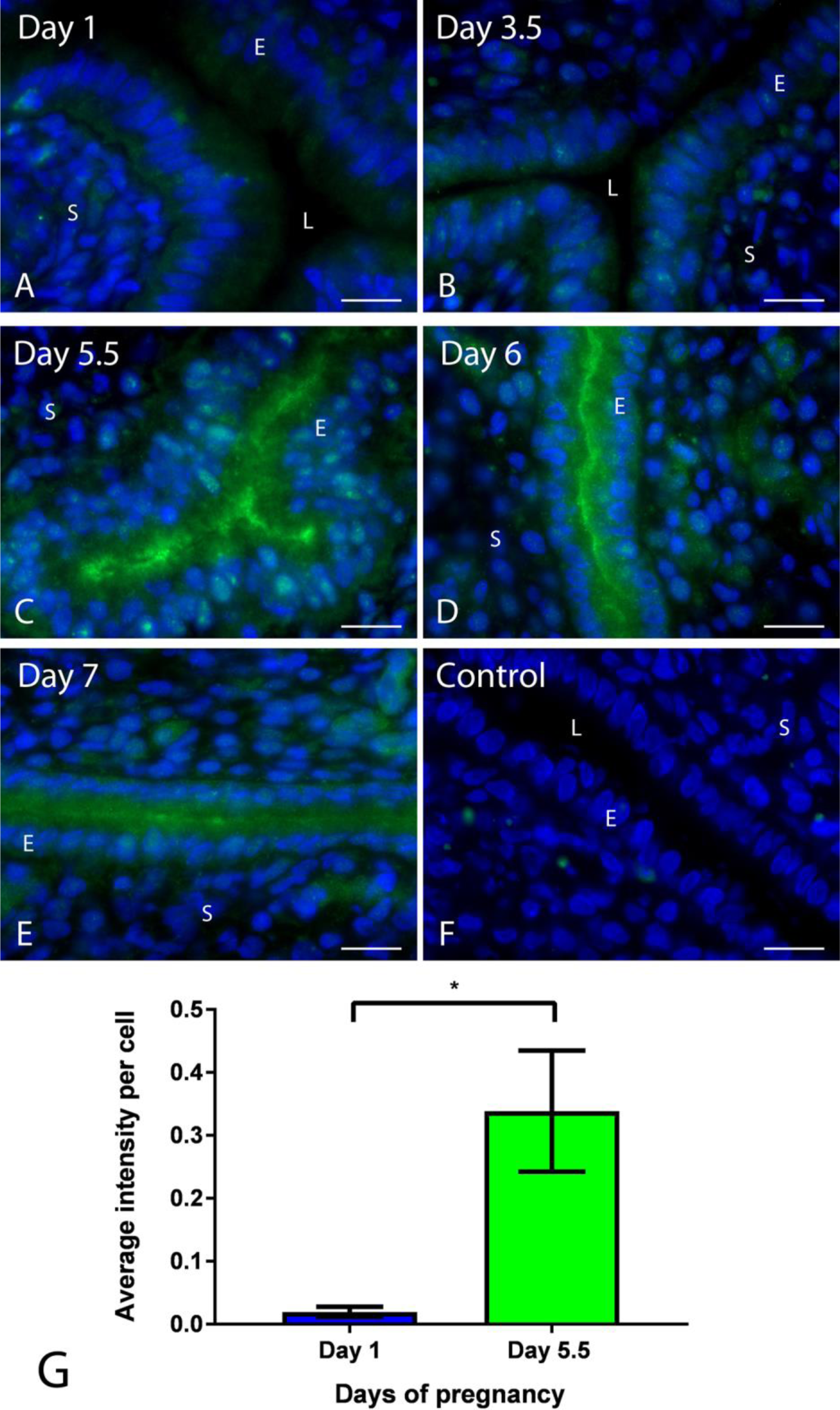
Syntaxin 2 localisation in rat uterus (A, B) Day 1 and 3.5 of pregnancy shows diffuse cytoplasmic localisation of syntaxin 2 in UECs. (C, D) Day 5.5 and 6 of pregnancy syntaxin 2 is concentrated in the apical region of UECs. (E) Day 7 of pregnancy syntaxin 2 is apically cytoplasmic. (F) Non-immune control shows no staining in UECs. (G) Unpaired two-tailed Student’s t-test found syntaxin 2 had a significantly greater intensity in day 5.5 compared to day 1 of pregnancy in UECs. (* P<0.05) n=5; error bar is the mean ± S.E.M. All scale bars are 20µm. (L= lumen, E= epithelium and S= stroma)

### SNAP23 is present in UECs during early pregnancy

SNAP23 is present in UECs during early pregnancy in the rat uterus when examined through indirect immunofluorescence microscopy (Fig. 4). On days 1, 3.5, 6 and 7, SNAP23 is localised throughout the cytoplasm in UECs (Fig. 4A, B, D, E). On day 5.5 however, SNAP23 is localised apically (Fig. 4C). Non-immune controls were performed with all immunofluorescence protocols showing no staining in UECs. A representative image of day 5.5 non-immune controls is shown (Fig. 4F). Fluorescent intensity measurements found significantly more SNAP23 is staining per epithelial cell on day 5.5 compared to day 1 of pregnancy; *p <0.05; n=5 (Fig. 4G).

**Figure 4:**
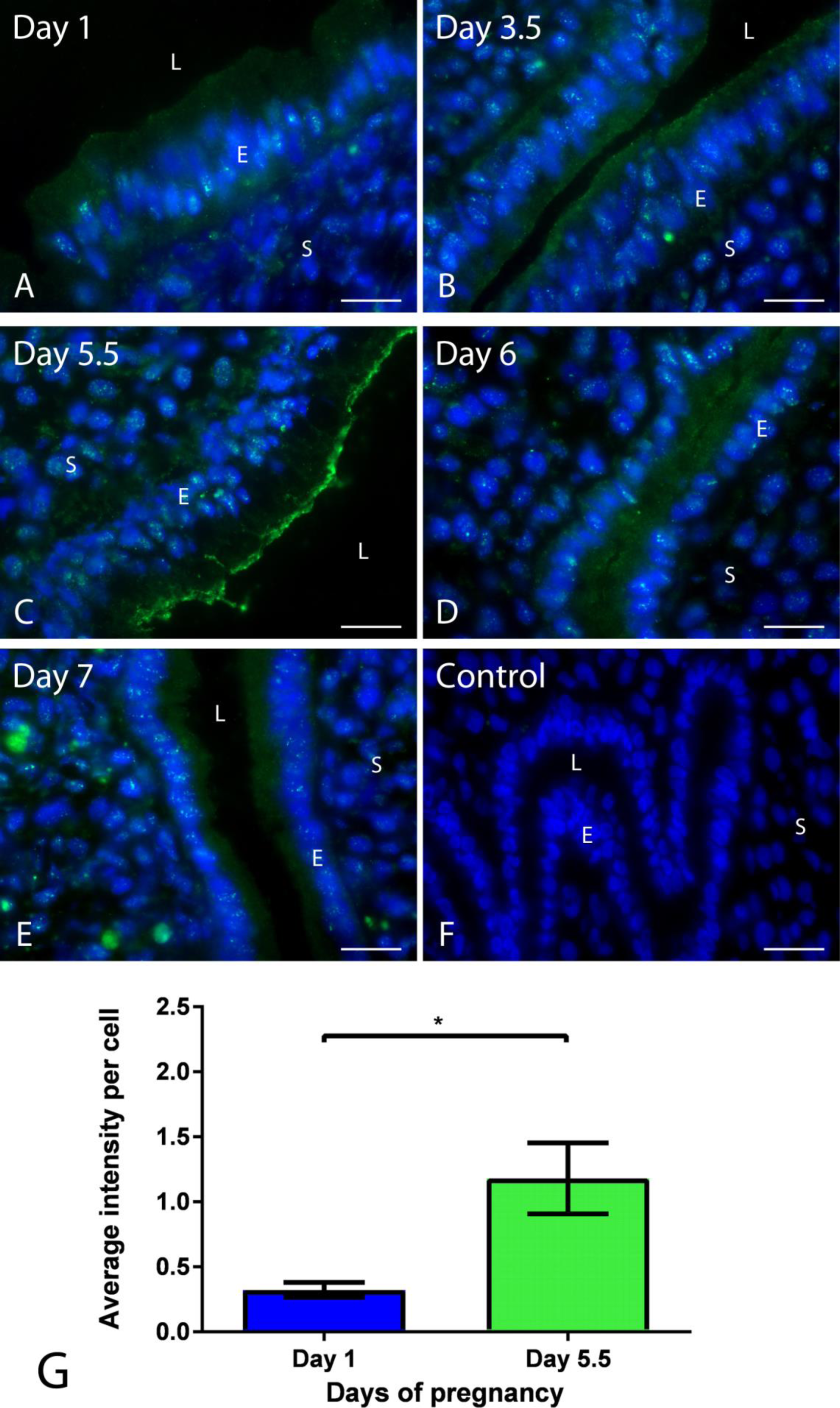
SNAP23 localisation in rat uterus (A, B) Day 1 and 3.5 of pregnancy shows diffuse cytoplasmic localisation of SNAP23 in UECs. (C) Day 5.5 of pregnancy SNAP23 is concentrated in the apical region of UECs. (D, E) Day 6 and 7 of pregnancy SNAP23 is cytoplasmic. (F) Non-immune control shows no staining in UECs. (G) Unpaired two-tailed Student’s t-test found SNAP23 had a significantly greater intensity in day 5.5 compared to day 1 of pregnancy in UECs. (* P<0.05) n=5; error bar is the mean ± S.E.M. All scale bars are 20µm. (L= lumen, E= epithelium and S= stroma)

### Western blot analysis of syntaxin 2 and SNAP23

Western blot analysis found syntaxin 2 (33 kDa) present in UECs on all days of early pregnancy examined and that the amount of syntaxin 2 was significantly higher on day 5.5 compared to day 1 (Fig. 5A, C). Similarly, western blot analysis found SNAP23 (57 kDa) in UECs on all days of early pregnancy (Fig. 5B), but that SNAP23 was significantly higher on day 5.5 compared to day 1 and 7. A significant decrease in SNAP23 was also observed between day 3.5 and 7 (Fig. 5D).

**Figure 5:**
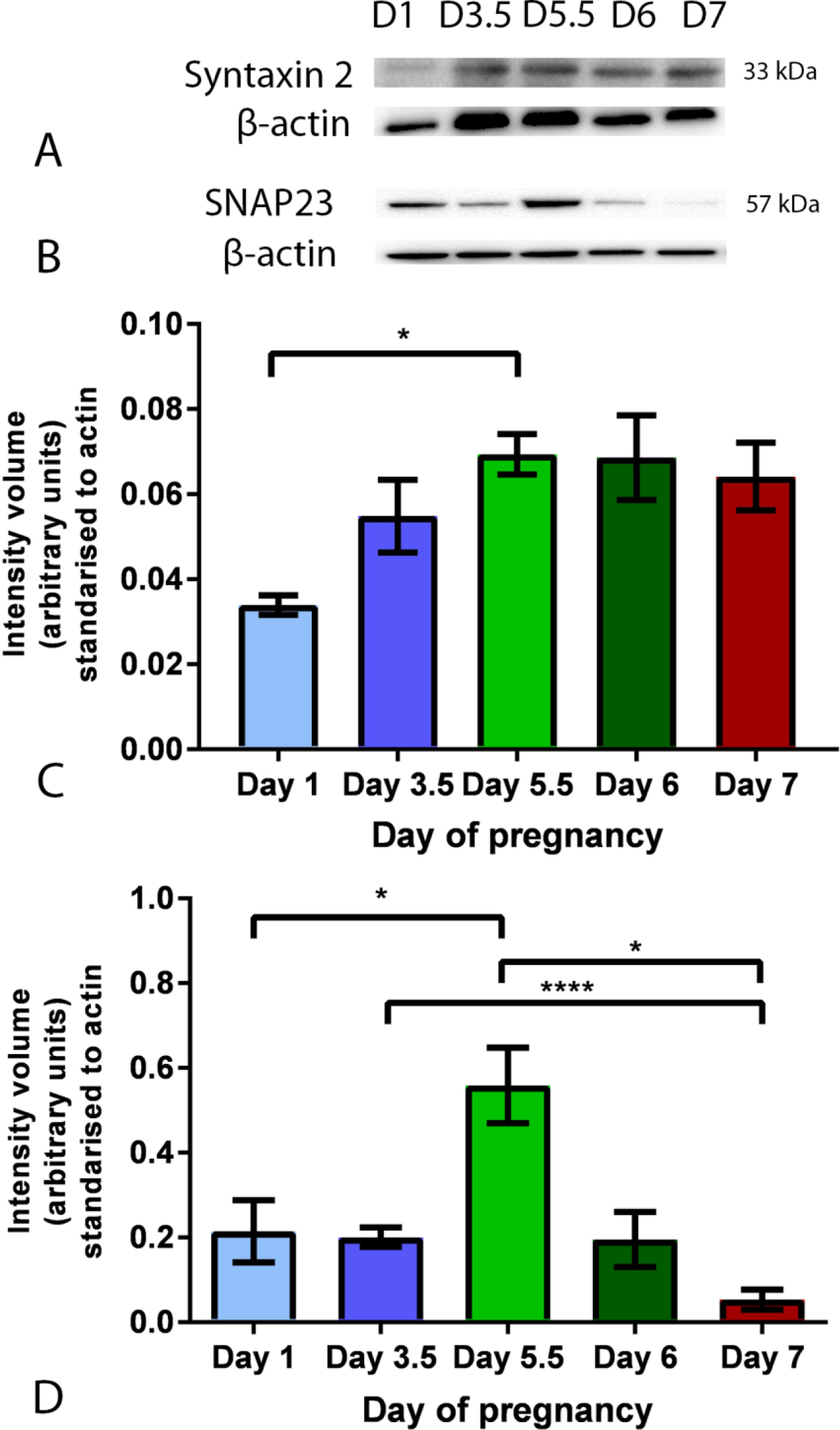
Western blot analysis of syntaxin 2 and SNAP23 in isolated UECs (A) Syntaxin 2 is present at 33 kDa in isolated UECs during day 1, 3.5, 5.5, 6 and 7 of pregnancy. (B) SNAP23 is present at 57 kDa in isolated UECs during day 1, 3.5, 5.5, 6 and 7 of pregnancy. β-actin was used as a loading control. (C) Densitometric and statistical analysis (one-way ANOVA) found a significant increase in syntaxin 2 in day 5.5 compared to day 1 (*P<0.05) n=5. (D) Densitometric and statistical analysis (one-way ANOVA) found a significant increase in SNAP23 in day 5.5 compared to day 1. SNAP23 also showed a significantly decrease in day 7 compared to day 3.5 and 5.5 (*P<0.05, ***P<0.0001) n=5. Error bar is the mean ± S.E.M.

### SNAP23 is present in uterine luminal secretion at day 5.5 of pregnancy

Co-localisation experiments between SNAP23 and Phalloidin in UECs on day 5.5 revealed no apparent co-localisation of SNAP23 and actin (Fig. 6). SNAP23 staining was localised in the entire cavity of the luminal space on day 5.5 (Fig. 6A, B). Phalloidin stained apically and at the apico-lateral junctional areas on day 5.5 of pregnancy in UECs, representing the location of actin cytoskeleton and apical border of UECs (Fig. 6A, C). PCC was calculated and found an average PCC of 0.117; n=5 (S Fig. 3A). SNAP23 only (S Fig. 3B) and Phalloidin only (S Fig. 3C) control was also imaged with all three filters and the same parameters and found there was no cross talk or bleed through between the channels. Non-immune controls were conducted as described above and exhibited no staining (S Fig. 3D).

**Figure 6:**
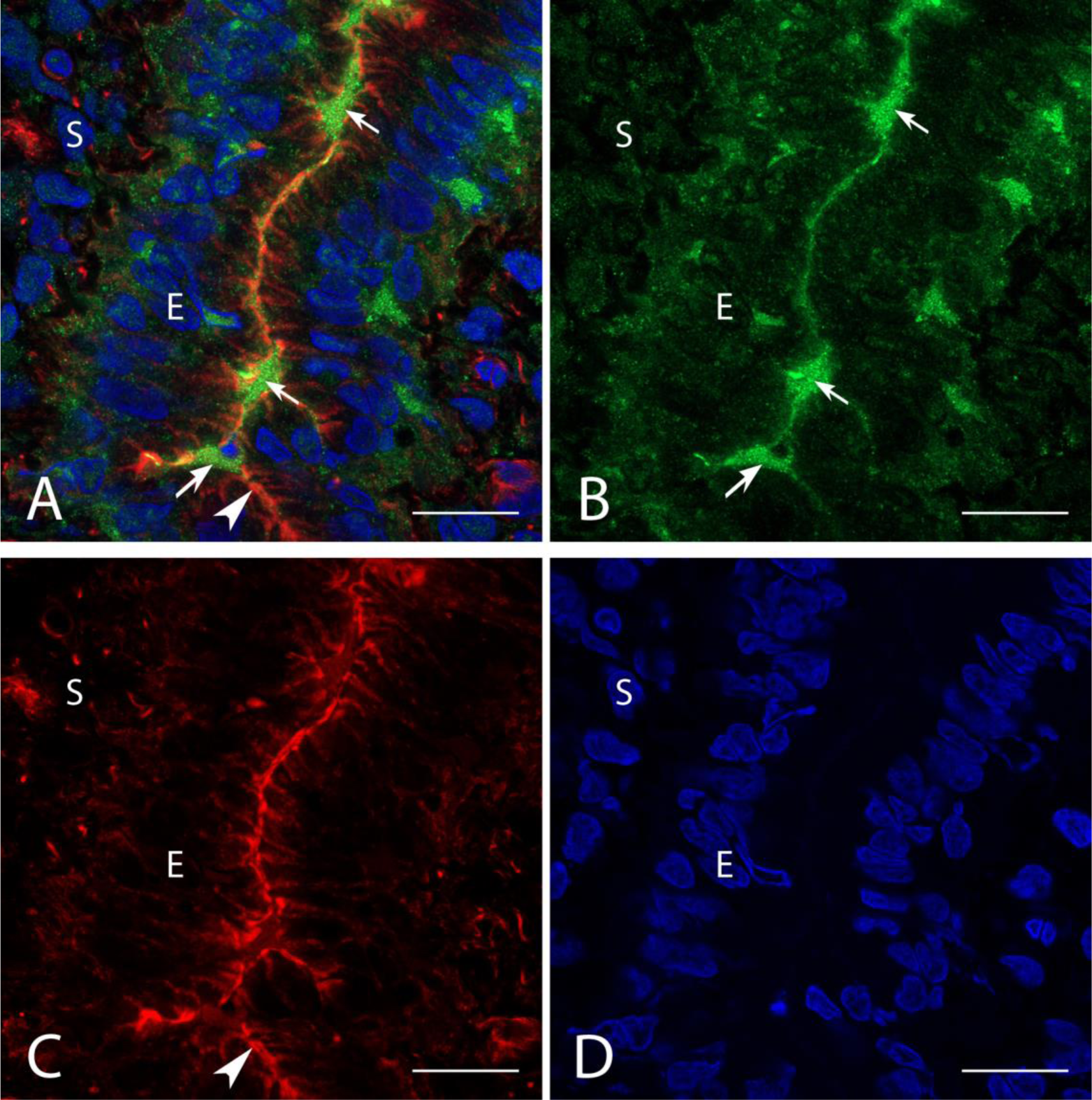
Confocal Microscopy-Triple labelling of SNAP23 and Phalloidin in UECs in day 5.5 of pregnancy. Merged channels showing the localisation of Phalloidin, SNAP23 and nuclei in UECs on day 5.5 of pregnancy. (B) Green channel shows SNAP23 (arrow) in present is secretions within the luminal space, outside the UECs. (C) Red channel shows Phalloidin (arrowhead) localises apically in UECs. (D) Blue channel shows the nuclei of UECs. All scale bars are 20µm. (L= lumen, E= epithelium and S= stroma)

## Discussion

This study is the first to detect that SNAP23 is present in the uterine luminal secretions exclusively during the time of receptivity in the rat. This indicates potential for SNAP23 to be used as a biomarker of a receptive endometrium, which could improve implantation rates during IVF protocols. The presence of syntaxin 2 and SNAP23 in UECs during early pregnancy, indicates that these t-SNAREs are both involved in the control of exocytosis and secretions that are known to occur at this time (Abonyo et al., 2004). This investigation also found a variety of EVs present in ULF. We suggest that SNAP23, syntaxin 2 and EVs collectively contribute to the microenvironment for blastocyst implantation.

Western blot data revealed that SNAP23 in UECs is found at 57 kDa on all days of early pregnancy and not at 23 kDa, as previously documented (Zhu et al., 2017). It is suggested that, in our study, SNAP23 (23 kDa) form a heterodimer with syntaxin 2 (34 kDa) resulting in a 57 kDa band. This higher band has been observed in other studies where syntaxin 2 and SNAP23 can form a heterodimer by interacting with VAMP2 to form a SNARE complex (Chen et al., 2000a; Chintagari et al., 2006; Hu et al., 2002). VAMP2 is in UECs during early pregnancy and is significantly higher at the time of receptivity (Kalam et al., 2020). This heterodimer, also referred to as an acceptor SNARE complex allows continued cycling of SNARE components for additional fusion events making exocytosis more efficient (Hong, 2005; Jahn and Scheller, 2006). The physical bond for SNAP and syntaxin heterodimers can be so strong that they are toxin and SDS-resistant (Low et al., 1998; Rao et al., 2004). We validated the SNAP23 antibody by confirming that the 23 kDa band is detected in non-polarised uterine endometrial immortalised cell lines HEC1A or RL95-2 cells (Sup Fig 2). Thus, it is expected that in highly polarised UECs as seen *in vivo*, SNAP23 exists as a heterodimer that plays a major role in the timely apical secretion of ULF responsible for maintaining the optimal microenvironment for blastocyst implantation.

SNAP23 mediates cytokine secretions in mast cells and synovial sarcoma cell lines (SW982 cells) (Boddul et al., 2014; Guo et al., 1998; Hepp et al., 2005; Suzuki and Verma, 2008). Cytokines IL-1β, TNFα and IL-6 are commonly found in human ULF collected during IVF procedures prior to embryo transfer (Boomsma et al., 2009; Rahiminejad et al., 2015). Unchanged amounts of IL-1β and lower levels of TNFα are found in women with successful pregnancies versus women with failed pregnancies (Rahiminejad et al., 2015). Thus, phosphorylated SNAP23 could be working with IL-1β to induce the release of TNFα and IL- 6 into the uterine fluid. In the uterus these cytokines are thought to mediate pro-inflammatory responses to facilitate communication between the UECs and the blastocyst to establish successful pregnancy (Chaouat et al., 2007).

Cellular actin is known to interact with SNAP23 to facilitate exocytosis (Eitzen, 2003; Lang et al., 2000; Pendleton and Koffer, 2001). Thus, this study performed co-localisation analysis of SNAP23 and actin in UECs and found that actin did not interact with SNAP23 (PCC of 0.117). Instead SNAP23 was found exclusively in the luminal space of the receptive uterus, where it appears to be part of the ULF. SNAP23 has previously been localised to the extracellular surface of platelets and phosphorylated SNAP23 has been shown to enable the release of exosomes in tumour cells (Flaumenhaft et al., 2007; Wei et al., 2017). It is likely that during the secretion process, SNAP23 is exocytosed itself or bound to other components of the secretory pathway such as in the membrane of EVs, which is observed in uterine fluid during early pregnancy in the rat.

The presence of syntaxin 2 in the apical region of UECs on day 5.5, 6 and 7 suggests a role in exocytosis during blastocyst implantation. Previous studies have found syntaxin 2 in platelets, pancreatic acinar cells, parotid acinar cells and alveolar type II cells, where it is involved in vesicular release (Abonyo et al., 2004; Chen et al., 1999; Imai et al., 2003; Pickett et al., 2005). In addition, inhibition of syntaxin 2 in platelets caused a dramatic 90% reduction in dense core granule release and inhibition of secretory lysosomes (Chen et al., 1999; Chen et al., 2000b). Further to this, in pancreatic acinar cells syntaxin 2 is present apically, is known to direct primary fusion and is found to change localisation to permit the onset of secondary fusion that results in compound exocytosis (Pickett et al., 2005). Compound exocytosis is a specialised form of secretion, where vesicles undergo fusion with each other as well as the target plasma membrane to optimise secretion (Pickett and Edwardson, 2006). The secretion in the uterine lumen during receptivity is unique and is produced for a finite period to create a microenvironment to assist implantation (Altmäe et al., 2017; Liang et al., 2017; Ng et al., 2013; Zhang et al., 2017). Our results indicate that in UECs, syntaxin 2 likely facilitating apical vesicular release and may promote be compound exocytosis to allow quicker release of vesicle content.

The EVs are a competent of the ULF that are secreted during early pregnancy and mediate intercellular communication between fetal and maternal tissue (Bidarimath et al., 2017).

EVs, exosomes and MVs were observed in the uterine lumen during early pregnancy and exhibited varying morphometric properties (namely size, diameter and membrane thickness). Exosomes, with two different membrane thickness were only seen on day 1 of pregnancy and not on day 5.5 or 6. Exosomes have previously been observed to be secreted by receptive ECC1 endometrial cells *in vitro* (Ng et al., 2013). However, in this study, exosomes were not seen to be secreted by rat UECs on the receptive days 5.5 and 6 of early pregnancy. This may be due to the lack of long branching glycocalyx chains at the time of receptivity (day 5.5 and 6) that are normally present on day 1 which may prevent these small exosomes being trapped on the surface of UECs in the same way they are on day 1 thus being washed away in TEM preparation (Murphy and Turner, 1991). Previous studies have found uterine exosomes during day 1 of pregnancy that fuse with the spermatozoa for sperm capacitation thus promoting their fertilising capacity (Franchi et al., 2016; Griffiths et al., 2008). The exosomes observed on day 1 in the present study may also be facilitating sperm capacitation.

This study showed MVs are present in the uterine lumen on day 1, 5.5 and 6 of pregnancy, with three different types of membranes with various thicknesses. Previous studies on vesicle membrane thickness have reported clathrin proteins which have heavy and light protein chains, can affect the membrane thickness of clathrin-coated vesicles. Furthermore, phosphorylation of membrane proteins in MVs secreted from prostate glands has been shown to increase membrane thickness (Kirchhausen et al., 2014; Ronquist and Brody, 1985). Membrane thickness is controlled by lipid-protein interactions and membrane protein coating that affects the distribution, organization, and function of bilayer-spanning proteins and cholesterol that in turn regulates membrane function such as membrane permeability, surface protein expression and free-energy changes (Andersen and Koeppe, 2007; Escribá et al., 2008; Kučerka et al., 2009; Paula et al., 1996). Therefore, membrane thickness is thought to differ depending on membrane cargo and MV function. It is therefore suggested here that these three different MVs observed in the ULF could be mediating cellular communication from the UECs towards the trophoblast cells to promote invasion into the uterus. As blastocysts injected with MVs isolated from the inner cell mass have better implantation outcomes compared to blastocyst injected with vehicle only in surrogate mice (Desrochers et al., 2016), the MVs secreted from the uterus found in ULF in our study may potentially be communicating and influencing embryo implantation and the establishment of pregnancy (Ng et al., 2013; Saadeldin et al., 2015; Tannetta et al., 2014).

These MVs were most obvious and seen to be secreted numerously from the UECs on day 5.5 of pregnancy, the time of receptivity. Here, type 1 and type 2 MVs were discovered to be larger in size compared to previous studies (Antonyak and Cerione, 2014; Ratajczak et al., 2006). This could be due to the fusion of multiple MVs with each other which has been observed in this study regardless of membrane type or size. The larger diameter and origin of secretion from non-apoptotic cells makes these MVs very similar to oncosomes (100nm- 10μm). Oncosomes are large EVs secreted by cancerous cells that facilitate transport of bioactive materials from one cell to another (Di Vizio et al., 2012; Morello et al., 2013). The different types of MVs may combine and exchange content resulting in MVs more suitable to communicate with and promote an invading blastocyst. MVs can have heterogenous membrane composition that carries multiple different proteins, lipids, nucleic acids and other bioactive molecules (Ratajczak et al., 2006). Thus, MVs are known to create a microenvironment that facilitates cell-cell communication and immune responses, which may be essential for promoting blastocyst implantation (Ratajczak et al., 2006; Tricarico et al., 2017).

This study investigated the control of EV secretion into the uterine luminal cavity during early pregnancy and showed that type 1 and 2 MVs are part of ULF secretions during early pregnancy. Both t-SNARES examined, syntaxin 2 and SNAP23, were found in UECs during early pregnancy and were significantly higher in abundance on day 5.5 (apposition) compared to day 1 of pregnancy. They were also present apically in UECs on day 5.5 compared to day 1. Overall, these finding show that SNAP23, syntaxin 2 and EVs may be collectively working together to create a favourable microenvironment for blastocyst implantation. This new knowledge about the composition and control of ULF could be used towards a non-invasive therapeutic approach in ART to determine time of receptivity in women leading up to embryo transfer to increase live birth rates through ART.

## Author contribution statement

S.N.K. designed the study, performed all experiments, analysed data and took the lead in writing the manuscript. All authors provided critical feedback and helped shape the research, analysis and manuscript.

## Acknowledgements

The authors acknowledge the scientific and technical assistance of the Bosch Institute Advanced Microscopy Facility, the Australian Microscopy and Microanalysis Research facility at the Australian Centre for Microscopy & Microanalysis, The University of Sydney. The author also thanks members of the Murphy lab.

## Supplementary

**S Figure 1:**
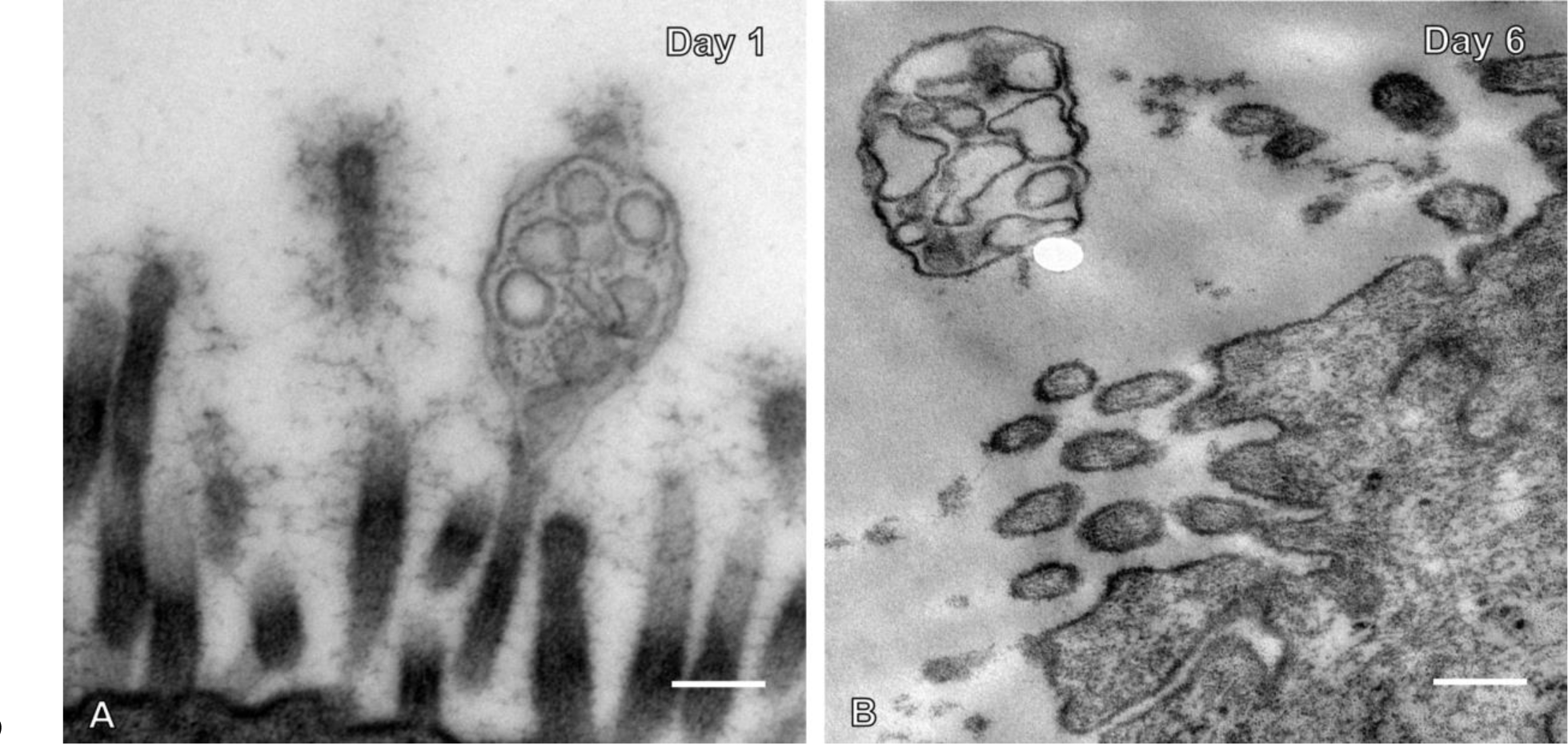
Transmission electron microscopy of multivesicular bodies on day 1 and 6 of early pregnancy. Multivesicular bodies were found in the pregnant uterine luminal space on day 1 (A) and day 6 (B). All scale bars are 200nm.

**S Figure 2:**
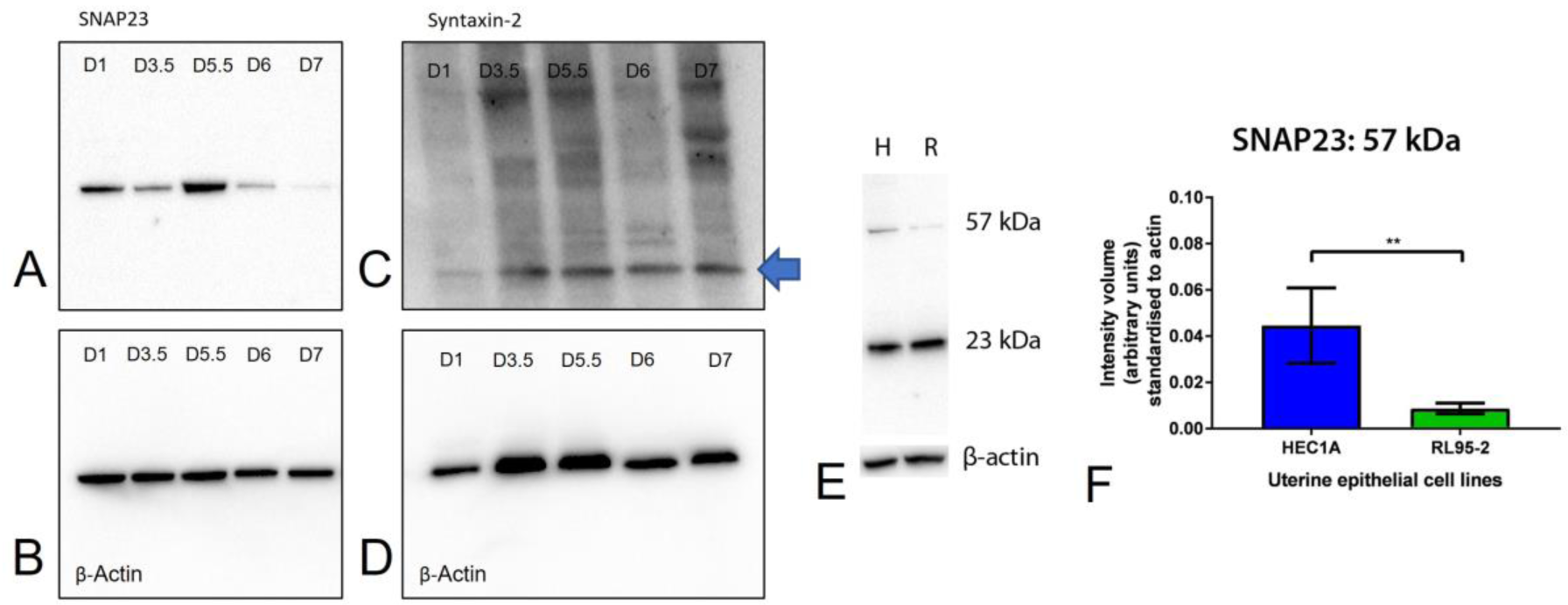
Blot transparency for SNAP23 and Syntaxin 2; and SNAP23 in UECs *in vitro* (A) Full, uncropped blot for SNAP23 at 57kDa and its corresponding β-actin blot (B). (C) Full, uncropped blot for Syntaxin 2 at 33 kDa (blue arrow) and corresponding β-actin blot (D). (E) SNAP23 is present at 23 kDa and 57 kDa in UECs *in vitro*. β-actin was used as a loading control. (F) Densitometry and statistical analysis (Student’s t-test) showed SNAP23 significantly decreased in RL95-2 compared to HEC1A cells, (** P<0.01) n=3.

**S Figure 3:**
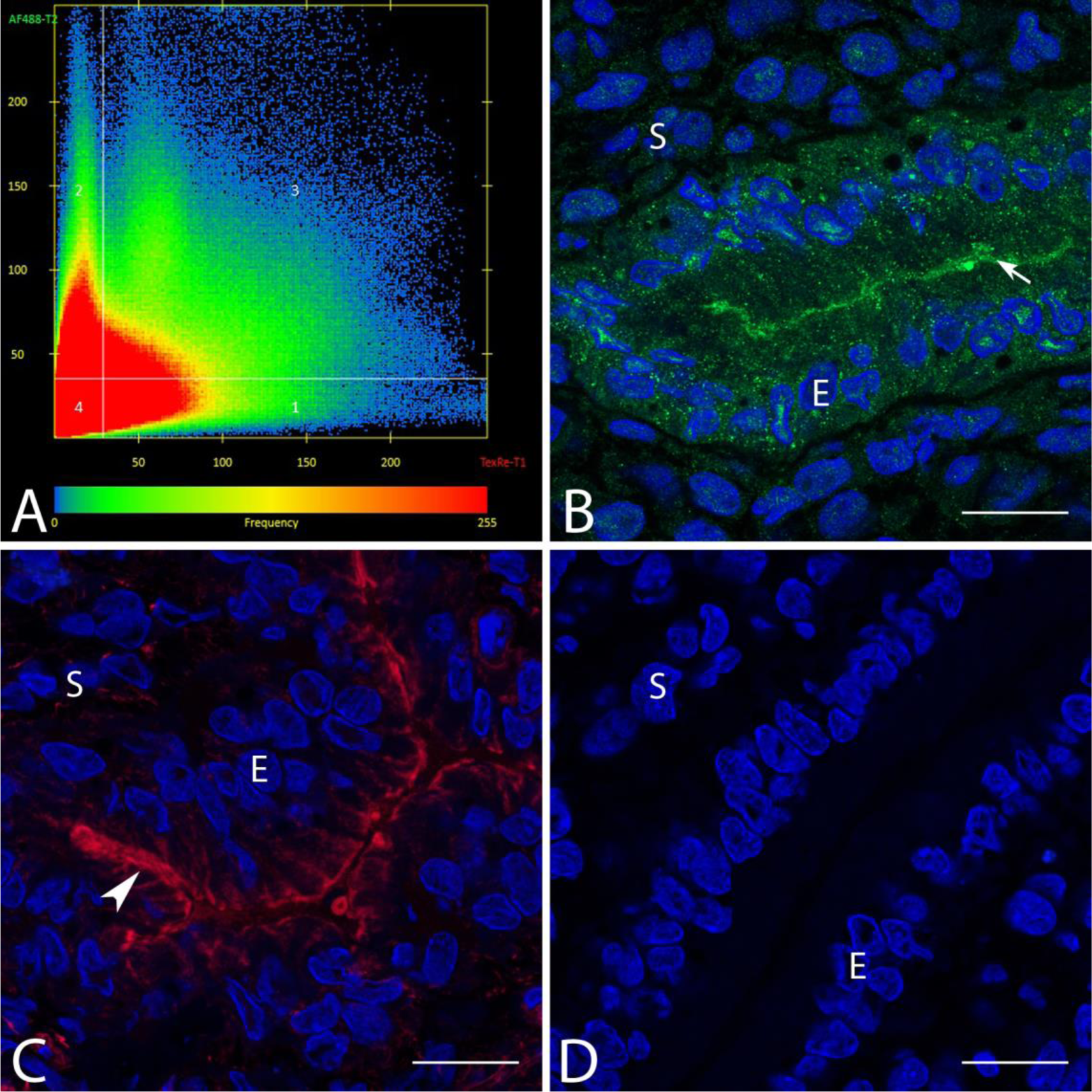
Confocal Microscopy-SNAP23 and Phalloidin co-localisation coefficient and controls (A) Pearson Co-localisation coefficient (PCC) scatter plot showing SNAP23 does not co- localise with Phalloidin, with an average PCC of 0.117, n=5. (B) SNAP23 staining (arrow) showing there is to cross talk (bleed through) from the red channel. (C) Phalloidin staining (arrowhead) showing there is to cross talk (bleed through) from the green channel. (D) Non- immune control shows no staining in UECs. All scale bars are 20µm. (L= lumen, E= epithelium and S= stroma)

## References

1. Abonyo, B. O., Gou, D., Wang, P., Narasaraju, T., Wang, Z. and Liu, L. (2004). Syntaxin 2 and SNAP-23 Are Required for Regulated Surfactant Secretion. Biochemistry 43, 3499–3506.

2. Aghajanova, L., Hamilton, A. E. and Giudice, L. C. (2008). Uterine receptivity to human embryonic implantation: Histology, biomarkers, and transcriptomics. Semin. Cell Dev. Biol. 19, 204–211.

3. Altmäe, S., Koel, M., Võsa, U., Adler, P., Suhorutšenko, M., Laisk-Podar, T., Kukushkina, V., Saare, M., Velthut-Meikas, A., Krjutškov, K., et al. (2017). Meta-signature of human endometrial receptivity: a meta-analysis and validation study of transcriptomic biomarkers. Sci. Rep. 7, 10077.

4. Andersen, O. S. and Koeppe, R. E. (2007). Bilayer Thickness and Membrane Protein Function: An Energetic Perspective. Annu. Rev. Biophys. Biomol. Struct. 36, 107–130.

5. Antonyak, M. A. and Cerione, R. A. (2014). Microvesicles as Mediators of Intercellular Communication in Cancer. In *Methods in molecular biology (Clifton*, N.J.), pp. 147– 173.

6. Aplin, J. D. (1997). Adhesion molecules in implantation. Rev. Reprod. 2, 84–93.

7. Bebelman, M. P., Smit, M. J., Pegtel, D. M. and Baglio, S. R. (2018). Biogenesis and function of extracellular vesicles in cancer. Pharmacol. Ther.

8. Bidarimath, M., Khalaj, K., Kridli, R. T., Kan, F. W. K., Koti, M. and Tayade, C. (2017). Extracellular vesicle mediated intercellular communication at the porcine maternal-fetal interface: A new paradigm for conceptus-endometrial cross-talk. Sci. Rep. 7, 40476.

9. Bobrie, A., Colombo, M., Raposo, G. and Théry, C. (2011). Exosome Secretion: Molecular Mechanisms and Roles in Immune Responses. Traffic 12, 1659–1668.

10. Boddul, S. V., Meng, J., Dolly, J. O. and Wang, J. (2014). SNAP-23 and VAMP-3 contribute to the release of IL-6 and TNFα from a human synovial sarcoma cell line. FEBS J. 281, 750–765.

11. Boomsma, C. M., Kavelaars, A., Eijkemans, M. J. C., Amarouchi, K., Teklenburg, G., Gutknecht, D., Fauser, B. J. C. M., Heijnen, C. J. and Macklon, N. S. (2009). Cytokine profiling in endometrial secretions: a non-invasive window on endometrial receptivity. Reprod. Biomed. Online 18, 85–94.

12. Burns, G., Brooks, K., Wildung, M., Navakanitworakul, R., Christenson, L. K. and Spencer, T. E. (2014). Extracellular Vesicles in Luminal Fluid of the Ovine Uterus. PLoS One 9, e90913.

13. Carnino, J. M., Lee, H. and Jin, Y. (2019). Isolation and characterization of extracellular vesicles from Broncho-Alveolar lavage fluid: A review and comparison of different methods. Respir. Res. 20,.

14. Carson, D. D., Bagchi, I., Dey, S. K., Enders, A. C., Fazleabas, A. T., Lessey, B. A. and Yoshinaga, K. (2000). Embryo Implantation. Dev. Biol. 223, 217–237.

15. Chaouat, G., Dubanchet, S. and Ledée, N. (2007). Cytokines: Important for implantation? J. Assist. Reprod. Genet. 24, 491–505.

16. Chen, D., Minger, S. L., Honer, W. G. and Whiteheart, S. W. (1999). Organization of the secretory machinery in the rodent brain: Distribution of the t-SNAREs, SNAP-25 and SNAP-23. Brain Res. 831, 11–24.

17. Chen, D., Bernstein, A. M., Lemons, P. P. and Whiteheart, S. W. (2000a). Molecular mechanisms of platelet exocytosis: role of SNAP-23 and syntaxin 2 in dense core granule release. Blood 95, 921–929.

18. Chen, D., Bernstein, A. M., Lemons, P. P. and Whiteheart, S. W. (2000b). Molecular mechanisms of platelet exocytosis: role of SNAP-23 and syntaxin 2 and 4 in lysosome release. Blood 95, 921–929.

19. Chintagari, N. R., Jin, N., Wang, P., Narasaraju, T. A., Chen, J. and Liu, L. (2006). Effect of Cholesterol Depletion on Exocytosis of Alveolar Type II Cells. Am. J. Respir. Cell Mol. Biol. 34, 677–687.

20. Cocucci, E., Racchetti, G. and Meldolesi, J. (2009). Shedding microvesicles: artefacts no more. Trends Cell Biol. 19, 43–51.

21. Crescitelli, R., Lässer, C., Szabó, T. G., Kittel, A., Eldh, M., Dianzani, I., Buzás, E. I. and Lötvall, J. (2013). Distinct RNA profiles in subpopulations of extracellular vesicles: apoptotic bodies, microvesicles and exosomes. J. Extracell. Vesicles 2, 20677.

22. Denzer, K., Kleijmeer, M. J., Heijnen, H. F., Stoorvogel, W. and Geuze, H. J. (2000). Exosome: from internal vesicle of the multivesicular body to intercellular signaling device. J. Cell Sci. 113 Pt 19, 3365–74.

23. Desrochers, L. M., Bordeleau, F., Reinhart-King, C. A., Cerione, R. A. and Antonyak, M. A. (2016). Microvesicles provide a mechanism for intercellular communication by embryonic stem cells during embryo implantation. Nat. Commun. 7, 11958.

24. Di Vizio, D., Morello, M., Dudley, A. C., Schow, P. W., Adam, R. M., Morley, S., Mulholland, D., Rotinen, M., Hager, M. H., Insabato, L., et al. (2012). Large oncosomes in human prostate cancer tissues and in the circulation of mice with metastatic disease. Am. J. Pathol. 181, 1573–84.

25. Eitzen, G. (2003). Actin remodeling to facilitate membrane fusion. Biochim. Biophys. Acta - Mol. Cell Res. 1641, 175–181.

26. EL Andaloussi, S., Mäger, I., Breakefield, X. O. and Wood, M. J. A. (2013). Extracellular vesicles: biology and emerging therapeutic opportunities. Nat. Rev. Drug Discov. 12, 347–357.

27. Erranz, B., Miquel, J. F., Argraves, W. S., Barth, J. L., Pimentel, F. and Marzolo, M.-P. (2004). Megalin and cubilin expression in gallbladder epithelium and regulation by bile acids. J. Lipid Res. 45, 2185–2198.

28. Escribá, P. V, González-Ros, J. M., Goñi, F. M., Kinnunen, P. K. J., Vigh, L., Sánchez-Magraner, L., Fernández, A. M., Busquets, X., Horváth, I. and Barceló-Coblijn, G. (2008). Membranes: a meeting point for lipids, proteins and therapies. J. Cell. Mol. Med. 12, 829–75.

29. Filant, J. and Spencer, T. E. (2014). Uterine glands: Biological roles in conceptus implantation, uterine receptivity and decidualization. Int. J. Dev. Biol. 58, 107–116.

30. Flaumenhaft, R., Rozenvayn, N., Feng, D. and Dvorak, A. M. (2007). SNAP-23 and syntaxin-2 localize to the extracellular surface of the platelet plasma membrane. Blood 110, 1492–1501.

31. Franchi, A., Cubilla, M., Guidobaldi, H. A., Bravo, A. A. and Giojalas, L. C. (2016). Uterosome-like vesicles prompt human sperm fertilizing capability. Mol. Hum. Reprod. 22, 833–841.

32. Glasser, S. R., Julian, J., Munir, M. I. and Soares, M. J. (1987). Biological markers during early pregnancy: trophoblastic signals of the peri-implantation period. Environ. Health Perspect. 74, 129–147.

33. Griffiths, G. S., Galileo, D. S., Reese, K. and Martin-Deleon, P. A. (2008). Investigating the role of murine epididymosomes and uterosomes in GPI-linked protein transfer to sperm using SPAM1 as a model. Mol. Reprod. Dev. 75, 1627–1636.

34. Guo, Z., Turner, C. and Castle, D. (1998). Relocation of the t-SNARE SNAP-23 from lamellipodia-like cell surface projections regulates compound exocytosis in mast cells. Cell 94, 537–548.

35. Hepp, R., Puri, N., Hohenstein, A. C., Crawford, G. L., Whiteheart, S. W. and Roche, P. A. (2005). Phosphorylation of SNAP-23 regulates exocytosis from mast cells. J. Biol. Chem. 280, 6610–6620.

36. Hong, W. (2005). SNAREs and traffic.

37. Hu, K., Carroll, J., Fedorovich, S., Rickman, C., Sukhodub, A. and Davletov, B. (2002). Vesicular restriction of synaptobrevin suggests a role for calcium in membrane fusion. Nature 415, 646–650.

38. Imai, A., Nashida, T., Yoshie, S. and Shimomura, H. (2003). Intracellular localisation of SNARE proteins in rat parotid acinar cells: SNARE complexes on the apical plasma membrane. Arch. Oral Biol. 48, 597–604.

39. Jahn, R. and Scheller, R. H. (2006). SNAREs — engines for membrane fusion. Nat. Rev. Mol. Cell Biol. 7, 631–643.

40. Jahn, R. and Südhof, T. C. (1999). Membrane Fusion and Exocytosis. Annu. Rev. Biochem. 68, 863–911.

41. Kalam, S. N., Dowland, S., Lindsay, L. and Murphy, C. R. (2018). Microtubules are reorganised and fragmented for uterine receptivity. Cell Tissue Res. 1–11.

42. Kalam, S. N., Cole, L., Lindsay, L. and Murphy, C. R. (2020). Membrane trafficking directed by VAMP2 and syntaxin 3 in uterine epithelial cells. Reproduction 1,.

43. Kalam, S. N., Dowland, S., Lindsay, L. and Murphy, C. R. (2022). Porosomes in uterine epithelial cells: ultrastructural identification and characterisation during early pregnancy. J. Morphol.

44. Kaneko, Y., Lindsay, L. A. and Murphy, C. R. (2008). Focal adhesions disassemble during early pregnancy in rat uterine epithelial cells. Reprod. Fertil. Dev. 20, 892–899.

45. Kirchhausen, T., Owen, D. and Harrison, S. C. (2014). Molecular structure, function, and dynamics of clathrin-mediated membrane traffic. Cold Spring Harb. Perspect. Biol. 6, a016725.

46. Kučerka, N., Nieh, M.-P., Pencer, J., Sachs, J. N. and Katsaras, J. (2009). What determines the thickness of a biological membrane. Gen. Physiol. Biophys 28, 117–125.

47. Lang, T., Wacker, I., Wunderlich, I., Rohrbach, A., Giese, G., Soldati, T. and Almers, W. (2000). Role of Actin Cortex in the Subplasmalemmal Transport of Secretory Granules in PC-12 Cells. Biophys. J. 78, 2863–2877.

48. Lessey, B. A. (2000). Endometrial receptivity and the window of implantation. Bailliere’s Best Pract. Res. Clin. Obstet. Gynaecol. 14, 775–788.

49. Liang, J., Wang, S. and Wang, Z. (2017). Role of microRNAs in embryo implantation. Reprod. Biol. Endocrinol. 15, 90.

50. Low, S. H., Roche, P. A., Anderson, H. A., van Ijzendoorn, S. C., Zhang, M., Mostov, K. E. and Weimbs, T. (1998). Targeting of SNAP-23 and SNAP-25 in polarized epithelial cells. J. Biol. Chem. 273, 3422–30.

51. Lu, S., Peng, H., Zhang, H., Zhang, L., Cao, Q., Li, R., Zhang, Y., Yan, L., Duan, E. and Qiao, J. (2013). Excessive Intrauterine Fluid Cause Aberrant Implantation and Pregnancy Outcome in Mice. PLoS One 8, e78446.

52. Malmersjö, S., Di Palma, S., Diao, J., Lai, Y., Pfuetzner, R. A., Wang, A. L., McMahon, M. A., Hayer, A., Porteus, M., Bodenmiller, B., et al. (2016). Phosphorylation of residues inside the SNARE complex suppresses secretory vesicle fusion. EMBO J. 35, 1810–21.

53. Marca, V. La and Fierabracci, A. (2017). Insights into the Diagnostic Potential of Extracellular Vesicles and Their miRNA Signature from Liquid Biopsy as Early Biomarkers of Diabetic Micro/Macrovascular Complications. Int. J. Mol. Sci. 18, 1974.

54. Moore, C. L., Cheng, D., Shami, G. J. and Murphy, C. R. (2016). Correlated light and electron microscopy observations of the uterine epithelial cell actin cytoskeleton using fluorescently labeled resin-embedded sections. Micron 84, 61–66.

55. Morello, M., Minciacchi, V., de Candia, P., Yang, J., Posadas, E., Kim, H., Griffiths, D., Bhowmick, N., Chung, L., Gandellini, P., et al. (2013). Large oncosomes mediate intercellular transfer of functional microRNA. Cell Cycle 12, 3526–3536.

56. Murphy, C. R. (1993). The Plasma Membrane of Uterine Epithelial Cells: Structure and Histochemistry. Prog. Histochem. Cytochem. 27, 1–66.

57. Murphy, C. R. (1994). Plasma membrane transformation: a common response of uterine epithelial cells during the peri-implantation period. Cell Biol. Int. 18, 1115–1128.

58. Murphy, C. R. (2004). Uterine receptivity and the plasma membrane transformation. Cell Res. 14, 259–267.

59. Murphy, C. R. and Dwarte, D. M. (1987). Increase in cholesterol in the apical plasma membrane of uterine epithelial cells during early pregnancy in the rat. Acta Anat 128, 76–79.

60. Murphy, C. R. and Martin, B. (1985). Cholesterol in the plasma membrane of uterine epithelial cells: a freeze-fracture cytochemical study with digitonin. J. Cell Sci. 78, 163– 172.

61. Murphy, C. R. and Turner, V. F. (1991). Glycocalyx carbohydrates of uterine epithelial cells increase during early pregnancy in the rat. J. Anat. 177, 109–115.

62. Ng, Y. H., Rome, S., Jalabert, A., Forterre, A., Singh, H., Hincks, C. L. and Salamonsen, L. A. (2013). Endometrial Exosomes/Microvesicles in the Uterine Microenvironment: A New Paradigm for Embryo-Endometrial Cross Talk at Implantation. PLoS One 8, e58502.

63. Nguyen, H. P. T., Simpson, R. J., Salamonsen, L. A. and Greening, D. W. (2016). Extracellular vesicles in the intrauterine environment: Challenges and potential functions. Biol. Reprod. 95,.

64. Nichols, B. J., Ungermann, C., Pelham, H. R. B., Wickner, W. T. and Haas, A. (1997). Homotypic vacuolar fusion mediated by t- and v-SNAREs. Nature 387, 199–202.

65. Paula, S., Volkov, A. G., Van Hoek, A. N., Haines, T. H. and Deamer, D. W. (1996). Permeation of protons, potassium ions, and small polar molecules through phospholipid bilayers as a function of membrane thickness. Biophys. J. 70, 339–348.

66. Pendleton, A. and Koffer, A. (2001). Effects of latrunculin reveal requirements for the actin cytoskeleton during secretion from mast cells. Cell Motil. Cytoskeleton 48, 37–51.

67. Pickett, J. A. and Edwardson, J. M. (2006). Compound Exocytosis: Mechanisms and Functional Significance. Traffic 7, 109–116.

68. Pickett, J. A., Thorn, P. and Edwardson, J. M. (2005). The plasma membrane Q-SNARE syntaxin 2 enters the zymogen granule membrane during exocytosis in the pancreatic acinar cell. J. Biol. Chem. 280, 1506–11.

69. Rahiminejad, M. E., Moaddab, A., Ebrahimi, M., Rabiee, S., Zamani, A., Ezzati, M. and Abdollah Shamshirsaz, A. (2015). The relationship between some endometrial secretion cytokines and in vitro fertilization. Iran. J. Reprod. Med. 13, 557–62.

70. Rao, S. K., Huynh, C., Proux-Gillardeaux, V., Galli, T. and Andrews, N. W. (2004). Identification of SNAREs involved in synaptotagmin VII-regulated lysosomal exocytosis. J. Biol. Chem. 279, 20471–9.

71. Raposo, G. and Stoorvogel, W. (2013). Extracellular vesicles: exosomes, microvesicles, and friends. J. Cell Biol. 200, 373–83.

72. Ratajczak, J., Wysoczynski, M., Hayek, F., Janowska-Wieczorek, A. and Ratajczak, M. Z. (2006). Membrane-derived microvesicles: important and underappreciated mediators of cell-to-cell communication. Leukemia 20, 1487–1495.

73. Ronquist, G. and Brody, I. (1985). The prostasome: its secretion and function in man. Biochim. Biophys. Acta 822, 203–18.

74. Saadeldin, I. M., Oh, H. J. and Lee, B. C. (2015). Embryonic–maternal cross-talk via exosomes: Potential implications. Stem Cells Cloning Adv. Appl. 8, 103–107.

75. Salamonsen, L. A., Evans, J., Nguyen, H. P. T. and Edgell, T. A. (2016). The Microenvironment of Human Implantation: Determinant of Reproductive Success. Am. J. Reprod. Immunol. 75, 218–225.

76. Singh, H. and Aplin, J. D. (2009). Adhesion molecules in endometrial epithelium: Tissue integrity and embryo implantation. J. Anat. 215, 3–13.

77. Söllner, T., Whiteheart, S. W., Brunner, M., Erdjument-Bromage, H., Geromanos, S., Tempst, P. and Rothman, J. E. (1993). SNAP receptors implicated in vesicle targeting and fusion. Nature 362, 318–324.

78. Sutton, R. B., Fasshauer, D., Jahn, R. and Brunger, A. T. (1998). Crystal structure of a SNARE complex involved in synaptic exocytosis at 2.4 A resolution. Nature 395, 347– 353.

79. Suzuki, K. and Verma, I. M. (2008). Phosphorylation of SNAP-23 by IκB Kinase 2 Regulates Mast Cell Degranulation. Cell 134, 485–495.

80. Tannetta, D., Dragovic, R., Alyahyaei, Z. and Southcombe, J. (2014). Extracellular vesicles and reproduction–promotion of successful pregnancy. Cell. Mol. Immunol. 11, 548–563.

81. Toner, J. P. and Adler, N. T. (1985). The role of uterine luminal fluid in uterine contractions, sperm transport and fertility of rats. J. Reprod. Fertil. 74, 295–302.

82. Tricarico, C., Clancy, J. and D’Souza-Schorey, C. (2017). Biology and biogenesis of shed microvesicles. Small GTPases 8, 220–232.

83. Wei, Y., Wang, D., Jin, F., Bian, Z., Li, L., Liang, H., Li, M., Shi, L., Pan, C., Zhu, D., et al. (2017). Pyruvate kinase type M2 promotes tumour cell exosome release via phosphorylating synaptosome-associated protein 23. Nat. Commun. 8,.

84. Zhang, Y., Chen, Q., Zhang, H., Wang, Q., Li, R., Jin, Y., Wang, H., Ma, T., Qiao, J. and Duan, E. (2015). Aquaporin-dependent excessive intrauterine fluid accumulation is a major contributor in hyper-estrogen induced aberrant embryo implantation. Cell Res. 25, 139–42.

85. Zhang, Y., Wang, Q., Wang, H. and Duan, E. (2017). Uterine Fluid in Pregnancy: A Biological and Clinical Outlook. Trends Mol. Med. 23, 604–614.

86. Zhu, J.-J., Liu, Y.-F., Zhang, Y.-P., Zhao, C.-R., Yao, W.-J., Li, Y.-S., Wang, K.-C., Huang, T.-S., Pang, W., Wang, X.-F., et al. (2017). VAMP3 and SNAP23 mediate the disturbed flow-induced endothelial microRNA secretion and smooth muscle hyperplasia. Proc. Natl. Acad. Sci. 201700561.

